# Male songbirds show higher coccidia oocyst burdens than females following anthelmintic treatment

**DOI:** 10.64898/2025.12.12.694047

**Authors:** KM Talbott, AM Tysver, SM Wanamaker, ED Ketterson

## Abstract

In wildlife research, wild-caught vertebrates are often given anti-parasitic drugs when brought into captivity to protect animal health and reduce confounding effects of parasitic infection on research outcomes. However, the impacts of antiparasitic drugs on non-target parasite taxa are understudied, especially regarding host sex. To help address this gap, we investigated the impact of an anthelmintic medication on a protozoan gut parasite by quantifying coccidia oocyst burden in the feces of wild-caught dark-eyed juncos (*Junco hyemalis*) before and after two oral doses of ivermectin. Prior to treatment, males showed higher oocyst burdens than females. Following treatment, ivermectin-treated males showed larger increases in oocyst burden compared to both control males and ivermectin-treated females, while there was no difference in baseline and post-treatment oocyst burden in treated females. Our results suggest that wild-caught songbirds should be housed separately by sex during ivermectin treatment and that male enclosures should be cleaned at a relatively higher frequency due to excessive oocyst shedding. It is unclear whether increases in oocyst burden among males were attributable to helminth removal or a direct impact of ivermectin on host immune function. Further work investigating sex differences in the impact of antiparasitic drugs on non-target parasite taxa is warranted.

## Introduction

Wild vertebrates can harbor an incredibly diverse array of micro- and macroparasites (Petney & Andrews, 1998; Poulin & Morand, 2000; Weinstein & Kuris, 2016), which poses a challenge for researchers looking to integrate wild-caught animals into captive colonies for research or conservation purposes. Within the host, one parasite taxon may regulate the abundance of other coinfecting taxa (Cox, 2001; Ferreira et al., 2025; Pedersen & Fenton, 2007), either positively (Bruchfeld et al., 2015; Lello et al., 2004) or negatively (Sassone-Corsi et al., 2016; Shen et al., 2019), leading to complex parasite community dynamics. In addition, the host’s immune system regulates parasite abundance, which may vary by parasite taxa (Schmid-Hempel, 2021). To protect colony health and reduce confounding effects of non-target parasites, researchers often administer antiparasitic drugs to wild-caught animals during a quarantine period before integration (Stringer & Linklater, 2014). However, administering a drug to reduce the abundance of one parasitic group may influence non-target parasites (Ferrari et al., 2009; Pedersen & Antonovics, 2013; Pedersen & Fenton, 2015). For example, experimentally treating wild rodents with ivermectin to reduce gastrointestinal nematodes increases both the prevalence and burden of coccidia, a gastrointestinal protozoan parasite (Knowles et al., 2013; Pedersen & Antonovics, 2013).

During quarantine, wild-caught birds are often treated with ivermectin to reduce gastrointestinal helminths and ectoparasites, and may also be treated with other drugs to reduce common bacterial pathogens, malarial blood parasites, and/or gastrointestinal coccidia. Coccidia outbreaks are especially hazardous in captive colonies, as stress and other factors can exacerbate natural coccidia infections and lead to host mortality (McGill et al., 2010). As a taxon, the coccidia include protozoan parasites of the genera *Isospora, Eimeria, Caryospora*, and *Lankastrella*, with *Isospora* considered the most common genus infecting passerines (Knight et al., 2018). Coccidia transmission occurs through the fecal-oral route; oocysts are shed via feces of infected hosts, sporulate under appropriate environmental conditions, and the resulting sporozoites are thereafter infective when ingested (Box, 1977; Knight et al., 2018).

Given the high prevalence of coccidial infections in wild birds (Boughton, 1937), the potential complications of these infections in captivity, and the prevalent use of ivermectin in wild-caught birds held in captivity, it is important to determine whether ivermectin increases the burden of coccidia in birds. We predicted that experimental ivermectin treatment would increase coccidia burden in wild-caught songbirds, as has been demonstrated in wild wood mice (*Apodemus sylvaticus*) (Knowles et al., 2013).

Because the impact of antiparasitic drugs on non-target parasites is understudied (Northover et al., 2018; Pedersen & Fenton, 2015), it is unclear whether such impacts vary by host demographics, such as sex. Sex differences in parasite prevalence and burden are commonly reported in wild birds (Dawson & Bortolotti, 2001; Robinson et al., 2008; Roulin et al., 2007; Stewart & Merrill, 2015; van Oers et al., 2010), which is thought to be attributable to sex differences in parasite exposure (Silveira & Calegaro-Marques, 2016; Valdebenito et al., 2024), endocrine dynamics (Folstad & Karter, 1992), and/or host immune function (Pap et al., 2010; Valdebenito et al., 2021). Importantly, there could also be sex differences in the impact of antiparasitic drugs on non-target parasites, for example through differences in impacts on immune function and/or pharmacokinetics. Historically, drug research and development has primarily employed male models, meaning that sex differences in the bioavailability, distribution, metabolism, and elimination of commonly used drugs may exist (Gandhi et al., 2004). Indeed, female mammals show higher levels of ivermectin in the plasma compared to males given the same dose (Lifschitz et al., 2006). If this pattern holds true in birds, we might expect the higher bioavailability of ivermectin in female birds to cause a larger increase in coccidia burden in females, relative to male birds. In addition, if ivermectin pharmacokinetics vary by sex, it is possible that side effects (*e.g*. weight loss) could also vary by sex. Importantly, if ivermectin has distinct effects on coccidia burden or mass, it would have important ramifications for best practices in treating captive birds with ivermectin.

In this study we investigated the relationship between ivermectin treatment and coccidia oocyst burden in wild-caught dark-eyed juncos (*Junco hyemalis hyemalis*) held in captivity. We quantified burden by counting coccidia oocysts in junco feces at baseline and after each of two dosing timepoints. Experimental birds received two oral doses of ivermectin (Carpenter & Harms, 2023), while control birds received two doses of a carrier control. We tested the hypotheses that 1) oocyst counts would vary between ivermectin-treated males and control males, and 2) the impact of ivermectin treatment on coccidia count would vary by sex. In addition, we tested for sex differences in the impact of treatment on host mass.

## Methods

### Animal Capture and Care

Migratory dark-eyed juncos were captured near Lake Monroe (Monroe County, Indiana) in March of 2022 using baited mist-nets. Captured juncos were transported to the nearby Kent Farm Biological Observatory (Monroe County, Indiana), where they were housed for the remainder of the experiment in temperature-controlled rooms (∼60°F). Each treatment (6–10 birds) was housed in a pair of interconnected 8” x 8” ft rooms. Bird sex was determined by PCR (*n* = 6 females and n = 17 males; see protocol in Supplement), and birds were housed in same-sex groups. During the experiment, birds were held under a constant photoperiod of 17L:7D (hrs light:dark); a relatively long photoperiod was selected to induce and maintain a reproductive state (Rowan, 1925). We used this approach to increase the probability of detecting sex differences in response to the ivermectin treatment that may be attributable to sex differences in immune function (Valdebenito et al., 2021). All birds were provided with ad libitum water and a diet of millet, sunflower seeds, and mealworms.

Most of the males of this study (16/17 males) were used in a prior experiment probing the impact of experimental *Plasmodium* inoculation on male junco sperm quality, using methods described previously (Talbott & Ketterson, 2023). Of the 16 males used for the *Plasmodium* inoculation experiment, nine were inoculated with *Plasmodium*-infected junco blood and the others were inoculated with uninfected junco blood. A period of approximately six weeks elapsed between experiments, which we predicted to be sufficient time for recovery from the acute phase of infection (Cornet et al., 2013; Sarquis-Adamson & MacDougall-Shackleton, 2016). Altogether, we found no significant impact of *Plasmodium* inoculation on coccidia oocyst dynamics or mass in junco hosts (see Supplement).

### Experimental manipulation and timeline

We collected fecal samples and mass data on three time points: 7 days before the first ivermectin dose (‘baseline’), 7 days after the first dose (‘post dose one’), and 7 days after the second dose (‘post dose two’); see Figure 1. On dosing days, experimental birds were given 0.03 mL of 0.2 mg/kg ivermectin (Noromectin, Norbrook Inc) diluted in 5% dextrose, while control birds were given 0.03 mL of 5% dextrose only (Carpenter & Harms, 2023); all doses were administered orally. Mass data were collected to the nearest 0.01 g using a digital scale. Fecal samples were collected by placing each bird in a wax paper-lined paper bag for 10 minutes and collecting feces into a sterile 1.5 mL Eppendorf tube. Samples were stored at 5°C for up to 24 hours before quantifying oocysts (oocysts/g feces) by using fecal float analysis (see Supplement for fecal float protocol). All fecal samples were collected between two to six hours before sunset, as the highest oocyst counts are expected during this time period (Brawner III & Hill, 1999; Hudman et al., 2000; López et al., 2007). Coccidia were identified as *Isospora*, as sporulated oocysts contained two sporocysts (Knight et al., 2018).

**Figure 1.**
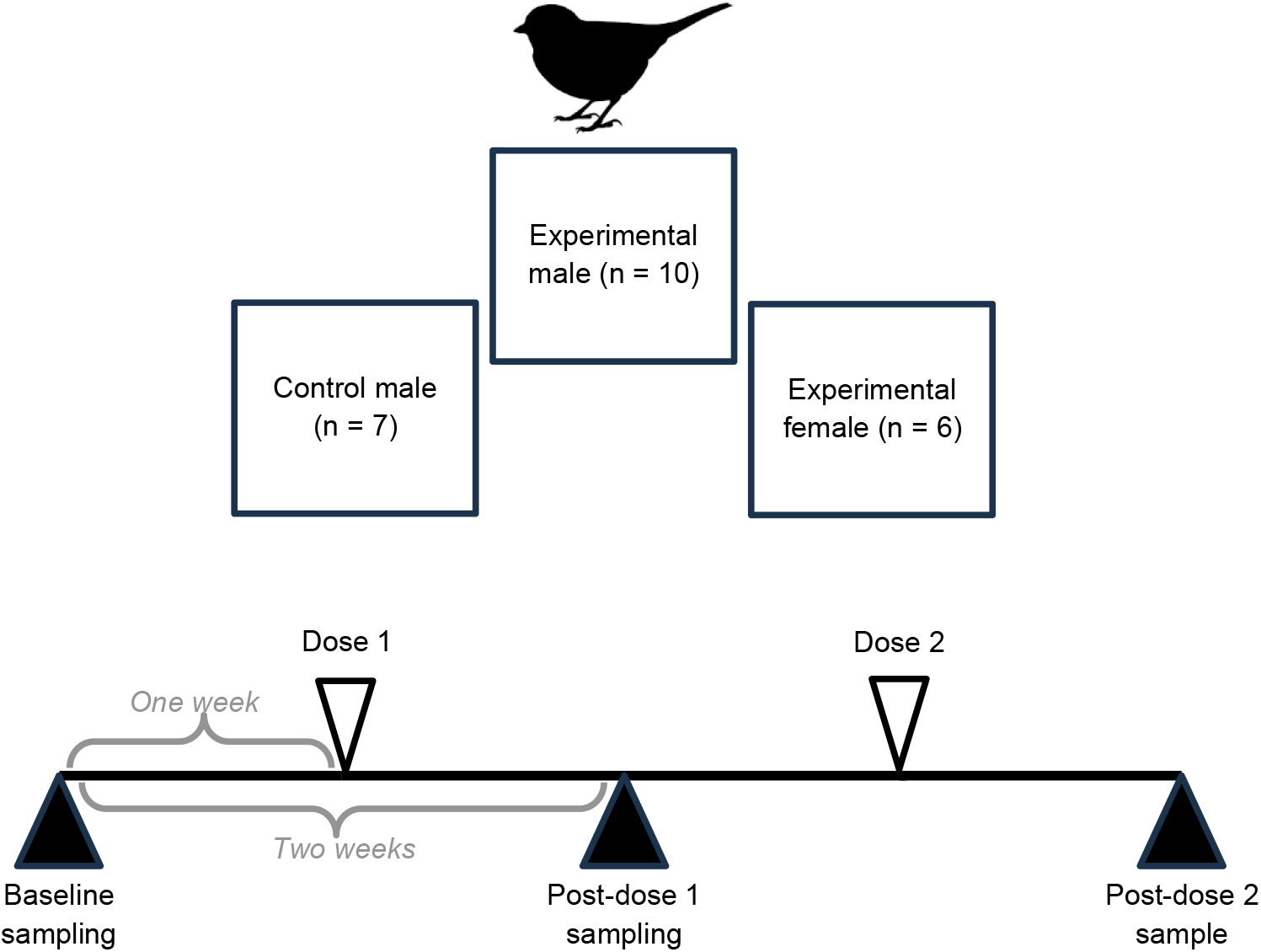
Experimental timeline and treatment groups. Feces and mass data were collected from dark-eyed juncos prior to manipulation (baseline), and then again at two and four weeks following baseline sampling. Treatment doses (ivermectin or carrier control) were administered at one week and three weeks following baseline sampling. Treatment groups included male and female juncos treated with ivermectin and one male control group given a carrier control.

### Statistical analyses

We used Wilcoxon rank-sum tests and Kruskal-Wallis tests for analyses of baseline oocyst counts and changes in oocyst counts from baseline to the two post-dose periods. For females only, (*n* = 6), we used a Wilcoxon signed-rank test to compare baseline to post-dose two oocyst counts.

Then, we used linear models to investigate predictors of mass change. We ran post-dose analyses using three data sets: ivermectin-treated birds, all males, and all data (with each bird assigned to one of three groups: ivermectin-treated males, ivermectin-treated females, or control males). All analyses were conducted in R version 2024.04.2.

## Results

### Baseline oocyst counts are higher in males than females

At baseline, oocyst counts were higher in males than in females (W = 82, *p* = 0.03; see Figure 2). Among males (*n* = 17), oocyst counts were higher in the ivermectin treatment group than in controls (*W* = 13, *p* = 0.03). Therefore, for subsequent analyses, we focused on changes in oocyst counts from baseline to post-dose (‘oocyst changes’), rather than raw oocyst counts.

**Figure 2.**
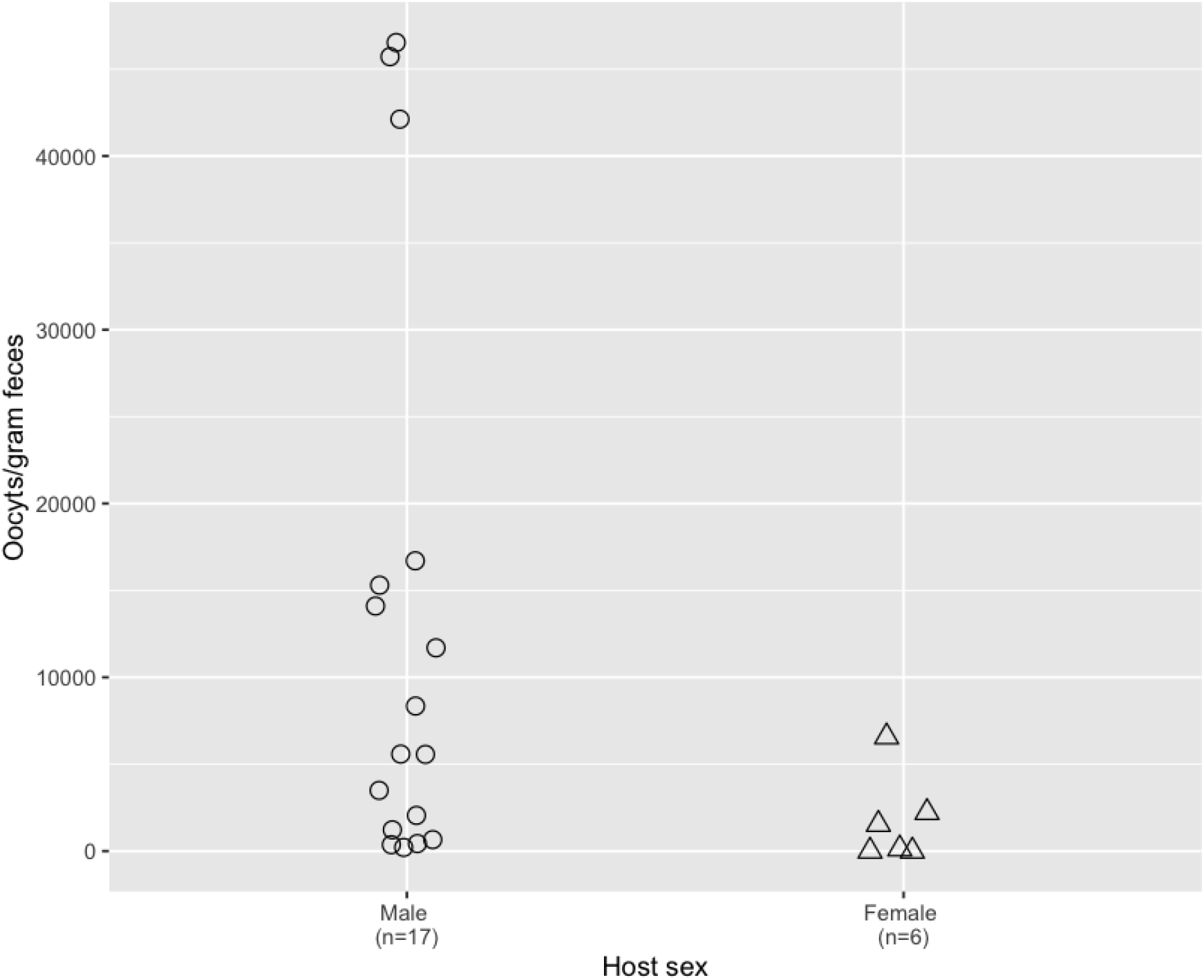
Pre-experiment (i.e., baseline) coccidia oocyst counts are higher in male dark-eyed juncos (*Junco hyemalis*) than in females. Points (jittered for easier viewing) represent data from individual juncos, with males shown in circles and females in triangles. Oocyst counts are quantified by oocysts per gram of feces.

### Ivermectin treatment increases oocyst counts after two doses

Among ivermectin-treated birds (*n* = 16), oocyst changes from baseline to post-dose one did not vary by host sex (*W* = 31, *p* = 0.96); see Figure 3A. Among all males (*n* = 17), oocyst changes did not vary by ivermectin treatment (*W* = 26, *p* = 0.42); see Figure 3B. Similarly, there were no differences in oocyst changes by group (*H*_*2*_ = 0.68, *p* = 0.71).

**Figure 3.**
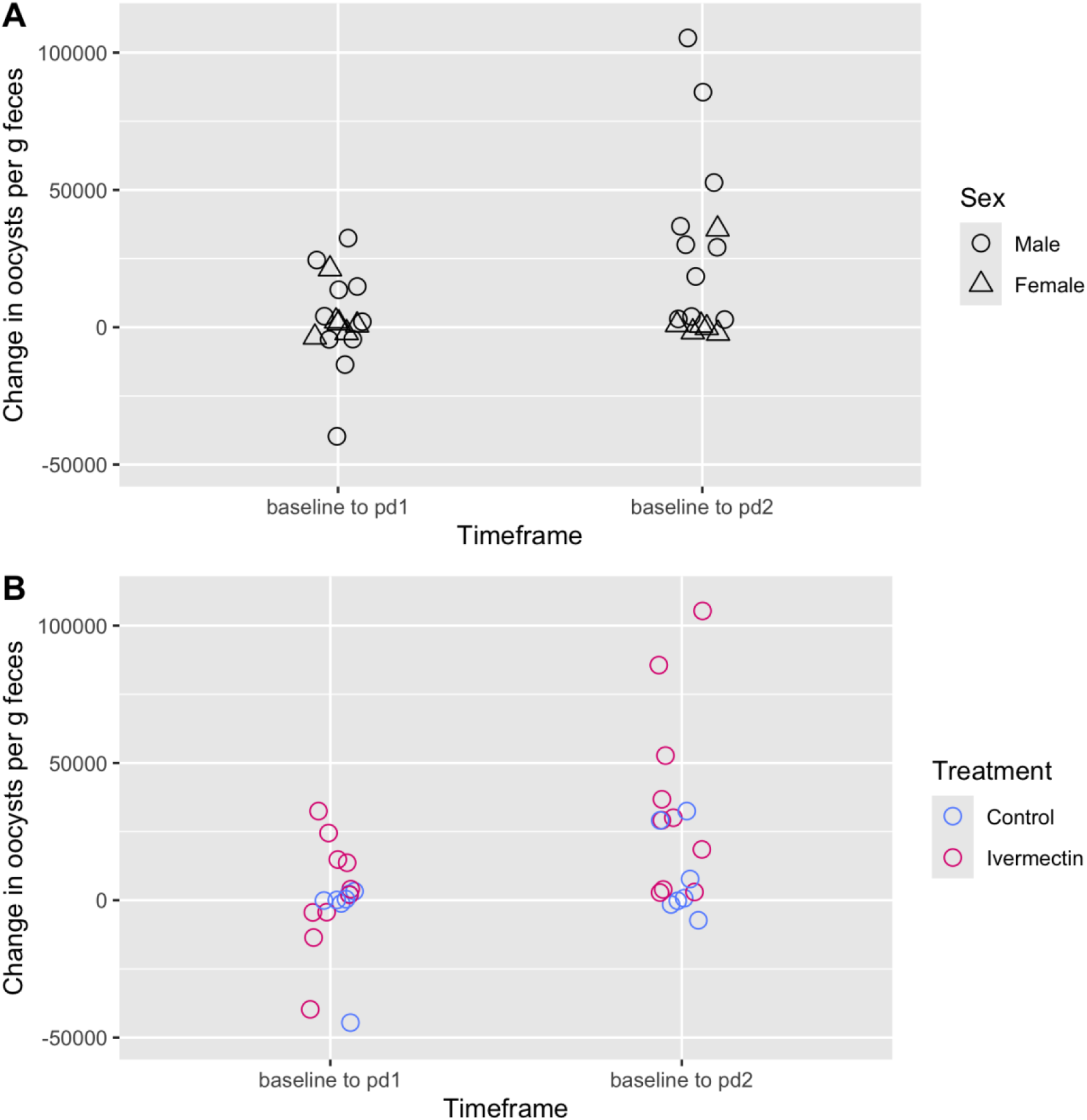
In dark-eyed juncos (*Junco hyemalis*), changes in coccidia oocyst counts (oocyst/g feces) are shown at two time points: after a first dose (pd1) and a second dose of ivermectin (pd2). A) Among ivermectin-treated juncos, coccidia oocyst counts do not vary by sex after the first dose, but males show larger increases from baseline than females following the second dose. B) Among all males, changes in oocyst count do not vary by ivermectin treatment following the first dose, but ivermectin treated males have larger increases in oocyst counts than control males after the second dose. Points represent data from individual juncos and are jittered for easier viewing.

Among ivermectin-treated birds, males showed larger, positive oocyst changes from baseline to post-dose two, compared to females (*W* = 54, *p* = 0.007); see Figure 3A. Among males, ivermectin-treated birds had larger, positive oocyst changes from baseline to post-dose two, compared to controls (*W* = 13, *p* = 0.03); see Figure 3B. Similarly, there was a difference in oocyst changes by group (*H*_*2*_ = 8.36, p = 0.02). A Dunn’s test with a Benjamini-Hochberg correction showed that both experimental females (*Z* = -2.19, adjusted *p* = 0.04) and control males (*Z* = -2.60, adjusted *p* = 0.03) had lower values than experimental males. Among females, there was no difference between baseline and post-dose two oocyst counts (*V* = 7, *p* = 1.00), suggesting there was no significant oocyst change within this group.

### No significant impact of treatment on host mass

Among ivermectin-treated birds, mass change from baseline to post-dose one did not vary by change in oocyst count (F_1,14_ = 0.02, Adj R^2^ < 0.01, *p* = 0.89) or sex (F_1,14_ = 0.01, Adj R^2^ < 0.01, *p* = 0.91); see Figure 4A. Among males, mass change did not vary by oocyst change (F_1,15_ = 0.02, Adj R^2^ < 0.01, *p* = 0.88), but there was a trend of greater mass loss in ivermectin-treated males compared to controls (Est = -0.80 ± 0.40, F_1,15_ = 3.97, Adj R^2^ = 0.16, *p* = 0.06). See Figure 4B. There were no differences in mass change by group (F_2,20_ = 1.20, Adj R^2^ = 0.02, *p* = 0.32).

**Figure 4.**
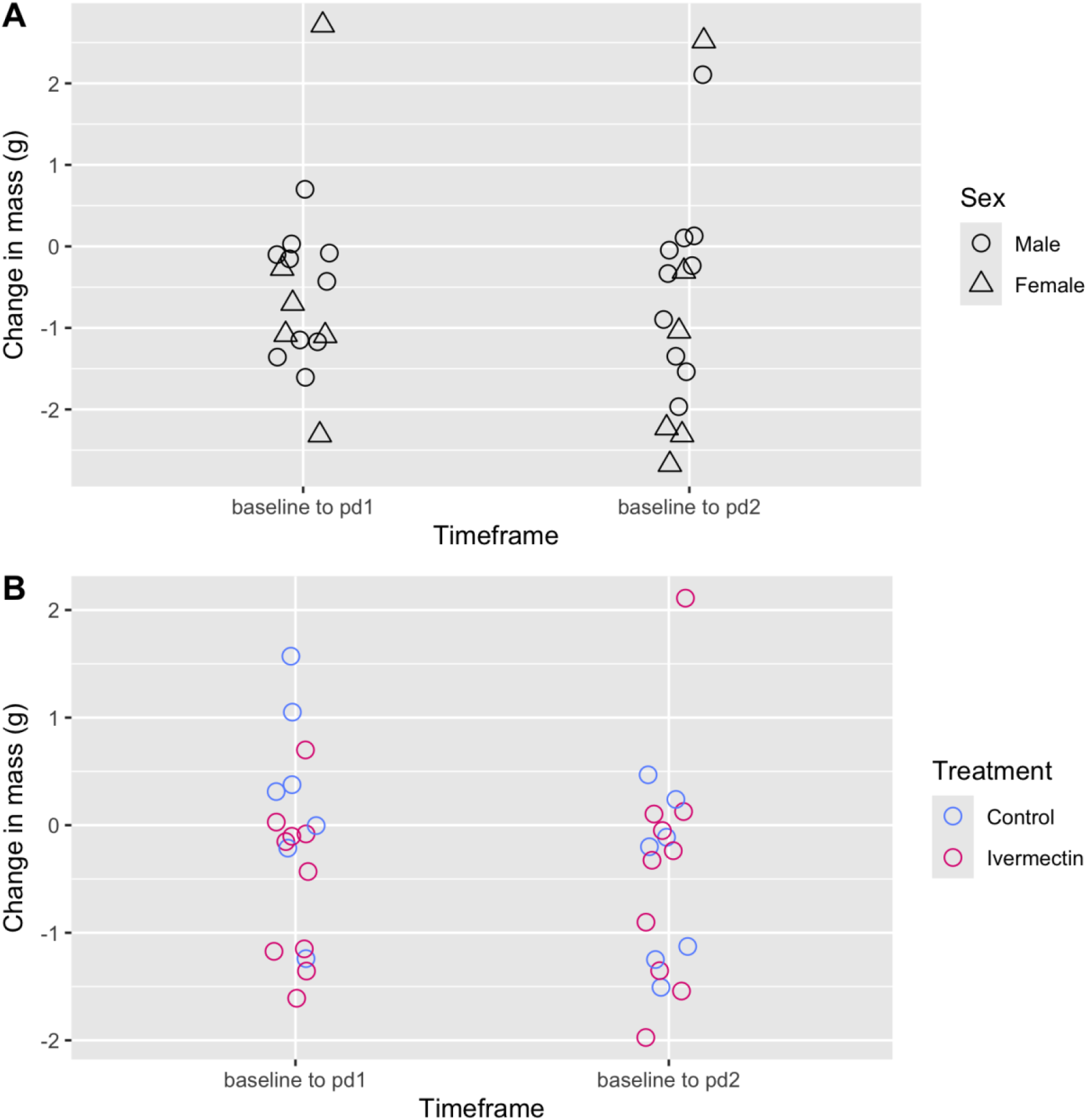
In dark-eyed juncos (*Junco hyemalis*), changes in mass (g) are shown at two time points: after a first dose (baseline to post-dose 1, ‘pd1’) and a second dose of ivermectin (baseline to post-dose 2, ‘pd2’). A) Among ivermectin-treated juncos, changes in mass to post-dose one and post-dose two did not vary by host sex. B) Among all males, there was a trend of higher mass loss among ivermectin-treated males compared to controls over the baseline to post-dose one timeframe. However, there were no treatment differences in mass changes from baseline to post-dose two. Points represent data from individual juncos and are jittered for easier viewing.

Among ivermectin-treated birds, mass changes from baseline to post-dose two did not vary by oocyst change (F_1,14_ = 0.41, Adj R^2^ < 0.1, *p* = 0.53) or host sex (F_1,14_ = 0.62, Adj R^2^ < 0.1, *p* = 0.44); see Figure 4A. Among males, mass changes did not vary by oocyst change (F_1,15_ = 1.06, Adj R^2^ < 0.1, p = 0.32) or ivermectin treatment (F_1,15_ = 0.04, Adj R^2^ < 0.1, *p* = 0.85). See Figures 4B. There were no differences in mass change by group (F_2,20_ = 0.42, Adj R^2^ < 0.01, *p* = 0.66).

## Discussion

Ivermectin is a commonly used anthelmintic drug in captive wildlife, yet its impacts on non-target parasite taxa in avian hosts remain unclear. In this study, we aimed to test the hypotheses that 1) ivermectin treatment would influence coccidia burden in male juncos and 2) ivermectin-treated juncos would vary in their coccidia burden by sex. We found support for both hypotheses, as larger post-treatment oocyst changes were found in ivermectin-treated males, compared to both control males and ivermectin-treated females, and there was no difference in between baseline and post-dose oocyst counts in ivermectin-treated females. This study provides insights into best practices for using ivermectin in wild-caught captive birds and suggests sex differences in the host-parasite dynamics of this system.

### Recommendations for the use of ivermectin in wild-caught songbirds

Our results indicate that ivermectin increases oocyst counts in wild-caught songbirds, at least in males. Therefore, we recommend cleaning bird enclosures and/or replacing bedding more frequently when treating captive passerines with ivermectin, especially enclosures containing male birds. We also recommend that researchers carefully weigh the costs and benefits of ivermectin treatment during quarantine; for example, if helminth prevalence is low, the benefits of mass ivermectin treatment during quarantine may be outweighed by the potential increase in coccidia burden, especially in group-housed birds. Furthermore, the male bias in oocyst burden both before and after ivermectin treatment may have negative impacts on female health when both sexes are co-housed. For example, if lower oocyst counts in females are driven by stronger immune regulation, the increased exposure to oocysts shed by co-housed males could exhaust females as they invest more resources in immune function. Even in the absence of ivermectin treatment, housing the sexes separately may be prudent, along with cleaning male enclosures at a higher frequency.

### Ivermectin and the ecoimmunology of avian gastrointestinal parasitism

In this study, we were unable to secure sufficient samples to determine pre-treatment helminth burden and identity. Therefore, it remains unclear whether the observed increase in coccidia burden is attributable to helminth removal or an immunomodulatory effect of ivermectin. A previous study showed increased coccidia burden in wild wood mice treated with ivermectin and suggested helminth removal as the most likely mechanism (Knowles et al., 2013). Because coccidia often co-occur in the host gut with helminths, the target parasite taxon for ivermectin, both parasite taxa may compete for similar resources, such as space and nutrients, while also being exposed the host’s immune system (Clerc et al., 2018; Lu et al., 2021; Saliba & Kirk, 2001). Coccidia live and reproduce within enterocytes lining the host’s gut (Lu et al., 2021) and helminth infection stimulates the host immune system to increase the rate of enterocyte turnover (Artis & Grencis, 2008). Thus, when the two taxa infect the same region of the gut, removing helminths may reduce the rate of enterocyte turnover, allowing coccidia infections to flourish (Knowles et al., 2013). In our study, birds were fed ad libitum, thereby reducing host-parasite competition for nutrients. Therefore, if helminth removal triggered the increase in coccidia burden in juncos, we also consider the most likely mechanism to be release from antagonistic effects of helminth infection on host immune function (Clerc et al., 2019; Knowles et al., 2013).

On the other hand, it is also possible that our results are directly attributable to immunomodulatory effects of ivermectin. For example, treating laboratory mice with ivermectin reduces T cell proliferation and the release of the pro-inflammatory cytokine INF-γ (Xie et al., 2023), which is an important component of the T helper style 1 host defense against coccidia development (Kim et al., 2019). There is also a sex difference in the impact of ivermectin on the in vitro production of TNF-ɑ, another pro-inflammatory cytokine, in wild wood mice; specifically, ivermectin treatment increased TNF-ɑ in male cell cultures, but reduced it in female cells (Rynkiewicz et al., 2019). While we would predict that this sex difference in TNF-ɑ production would yield decreases in oocyst burden in males and increases in oocyst burden in females, which is the opposite of what we observed, additional work investigating sex differences in the impact of ivermectin on songbird immune function is warranted.

### Sex differences in the impact of ivermectin on coccidia oocyst burden

We hypothesized that male and female juncos would differ in coccidia oocyst burden. This hypothesis was supported, as we found a male bias in both baseline oocyst counts and baseline to post-treatment oocyst changes. Similarly, naturally infected wild house finches show a male bias in oocyst burden when held in captivity (Weitzman et al., 2020). Further work is needed to determine whether this sex bias also occurs in passerines in a natural context, as several other studies found no sex differences in oocyst burden in avian hosts (Filipiak et al., 2009; Frigerio et al., 2016; Sykes et al., 2021) and one study showed a seasonal female bias in coccidia infection (Wascher et al., 2012). Interestingly, several studies in mammals have shown seasonally dependent male biases in oocyst burden, which tend to occur during periods of intense reproductive activity (Rijal et al., 2024; Strona et al., 2015) and/or at the end of the rainy season (Baines et al., 2015; Gorsich et al., 2014). Such seasonal effects suggest that oocyst shedding in vertebrates can be influenced by behaviors and/or physiological processes that vary by host sex. Thus, it is possible that the baseline male bias in oocyst burden we observed could be attributable to sex differences in immune regulation of coccidia, feeding behavior, and/or endocrinological responses to captivity.

Higher oocyst burdens in ivermectin-treated males compared to ivermectin-treated females suggests that the effects of ivermectin might vary by sex in birds. While relatively little is known about sex differences in the pharmacokinetics of ivermectin in birds, in most mammals, ivermectin reaches higher plasma concentrations and persists in plasma longer in females than males (Lifschitz et al., 2006; Ndong et al., 2007; Toutain et al., 1997); but see (Gokbulut et al., 2009). A female bias in ivermectin concentration was also shown locally in the gut epithelia and luminal contents of Wistar rats (Lifschitz et al., 2010). Possible explanations for this female bias include sex differences in body composition, endocrine dynamics, cellular structure of the gut, and feeding behavior, as food can bind the drug (Mariana et al., 2011; Ndong et al., 2007; Sartini et al., 2022). If these patterns hold true in birds, we would also expect higher bioavailability of ivermectin in the gut of female juncos, relative to males. Assuming similar helminth burdens between male and female juncos, we would therefore predict larger increases in oocyst burden in female juncos dosed with ivermectin, relative to males. However, we found the opposite pattern, with males showing a larger increase in oocyst burden following two doses of ivermectin, relative to females, and there was no change in oocyst burden in ivermectin-treated females. This would suggest that either 1) the bioavailability of ivermectin in orally dosed passerines is higher in males than females, counter to the pattern observed in most mammals, or 2) that pre-infection helminth burdens varied between the sexes. For example, if pre-treatment helminth burdens were higher in males than females, helminth removal might cause a larger increase in coccidia development in males than females. Additional work is needed to distinguish between these two possibilities.

## Acknowledgments

This project was funded by grants from the Wilson Ornithological Society, Indiana University Hutton Honors College, the Indiana University Research and Teaching Preserves. During this study, AMT was supported by the NSF Research Experience for Undergraduates through Indiana University’s Center for the Integrative Study of Animal Behavior (NSF award 2050311). We thank the summer 2022 Ketterson lab group for help with animal care and project support. Additionally, thank you to Laura Hurley and the summer 2022 CISAB REU program for mentorship and support of AMT throughout the project, and David Sinkiewicz in the IU CISAB Core Lab for use of equipment. Thank you to Randalyn Shepherd and Karen Rogers for assistance in developing the fecal floatation methods and providing guidance on ivermectin treatment protocols.

## Ethics statement

Birds were captured and banded under U.S. Fish and Wildlife Service Federal banding Permit #20261, Scientific Collecting Permit #MG093279-0, and Indiana Scientific Purposes License #3344. All methods were approved prior to research by the Indiana University Bloomington Institutional Animal Use and Care Committee (#21-029-3).

## Data availability statement

Data and code associated with this project are available at https://github.com/talbottkm/junco_ivermectin_coccidia

## Author contributions

All authors were involved in the conceptualization and design of the study. AT, KMT and SMW conducted the investigation. AT curated data. AT and KMT conducted formal analysis. KMT prepared visuals for the manuscript. KMT wrote the original draft and all authors contributed to review and editing of the final draft. SMW and EDK were responsible for project administration. EDK supervised the project.

## Supplement

### Effect of Plasmodium on oocyst burden in male juncos

We used Wilcoxon rank-sum tests to investigate differences in baseline oocyst counts, oocyst count changes, and mass between control and ivermectin-treated males. There was no difference in baseline oocyst count between males that had received a *Plasmodium* inoculation compared to those that did not (W = 37, p = 0.96, n = 17); see Figure S1. Among all males, changes in oocyst counts from baseline to post-dose one did not vary by *Plasmodium* treatment (W = 26, p = 0.37), nor did they vary from baseline to post-dose two (W = 38 = p = 0.89); see Figure S2. Male mass changes from baseline to post-dose one did not vary by *Plasmodium* treatment (W = 33, p = 0.81). Similarly, they did not vary from baseline to post-dose two (W = 33, p = 0.81); see Figure S3.

### Sexing PCR

We used PCR with primers P2 and P8 to determine junco sex (Griffiths et al., 1998). Each 10 μl reaction included 1.5 μl of each primer at 10 μm, 2 μl Go Taq Green Flexi Buffer (Promega), 0.2 μl Go Taq polymerase, 1 μl of dNTPs at 2 mM, 0.8 μl magnesium chloride at 25 mM, and 1 μl of template DNA. Products were run on 1% agarose gels stained with Gel Red (Biotium) at 90 V for 1 h and visualized under UV light. Birds scored as male later developed cloacal protuberances when held under a breeding photoperiod, confirming our sex assignments.

### Fecal float protocol

Prior to fecal float, weigh the sample by transferring it to a new 1.5 mL Eppendorf that has been tared on the scale. Weigh sample to the nearest 0.0001 g. Add one mL of 2.5% potassium dichromate solution to the Eppendorf tube containing the fecal sample, along with enough distilled water to bring the contents to a total volume of 1.5 mL. Centrifuge for five minutes at 1500 rpm; then, discard one mL of supernatant and add one mL of Sheather’s solution (Jorgenson Labs, Inc). Centrifuge again for five minutes at 1500 rpm. Drop additional Sheather’s solution into the tube to form a reverse meniscus and place a clean glass cover slip over the meniscus. Allow the sample to rest for 30 min while the oocysts float to the top and adhere to the cover slip. Then, count the number of oocysts observed under one field of view at 400x magnification on a compound light microscope. Divide the number of observed oocysts by pre-float sample mass.

### Figures

**Figure S1.**
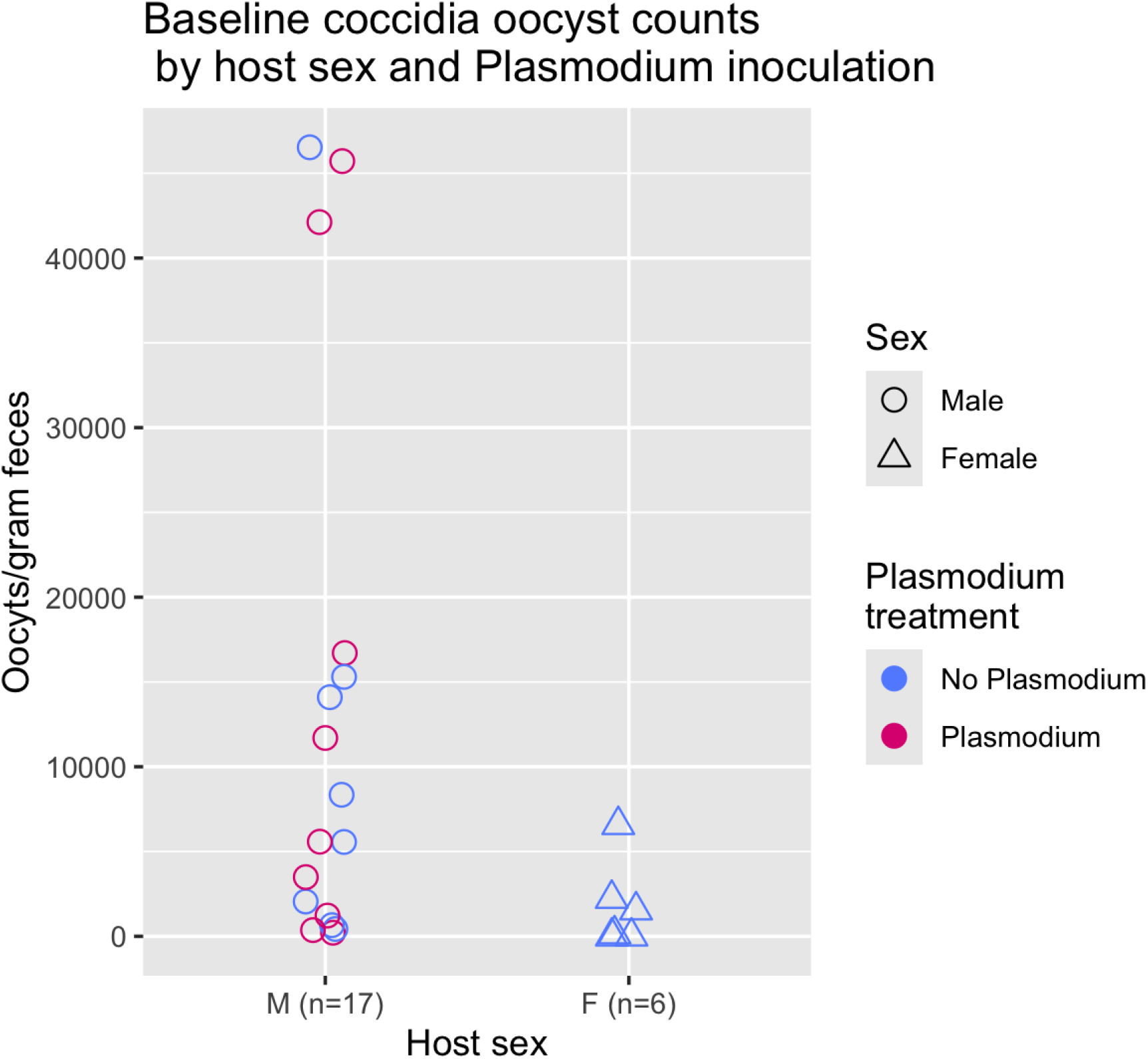
Baseline coccidia oocyst counts in dark-eyed juncos (*Junco hyemalis*) are higher in females than in males, but do not vary by *Plasmodium* treatment.

**Figure S2.**
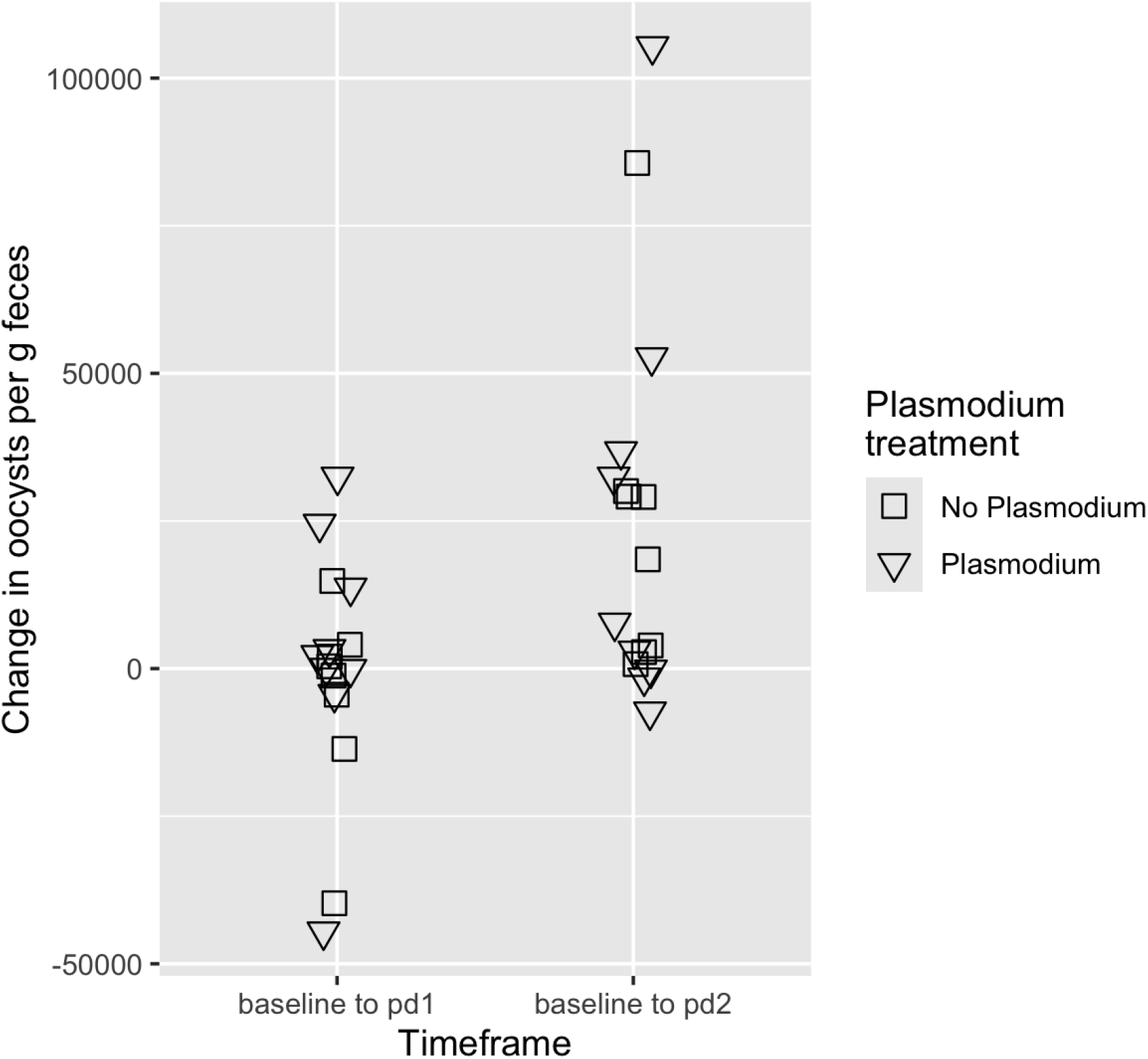
Changes in coccidia oocyst counts in male dark-eyed juncos (*Junco hyemalis*) do not vary by *Plasmodium* inoculation. Data represent changes in oocyst counts (oocyst/g feces) over two timeframes: from baseline to post-dose one, and baseline to post-dose two. At each dosing point, juncos were given either ivermectin or a carrier control. Points represent data from individual juncos and are jittered for easier viewing.

**Figure S3.**
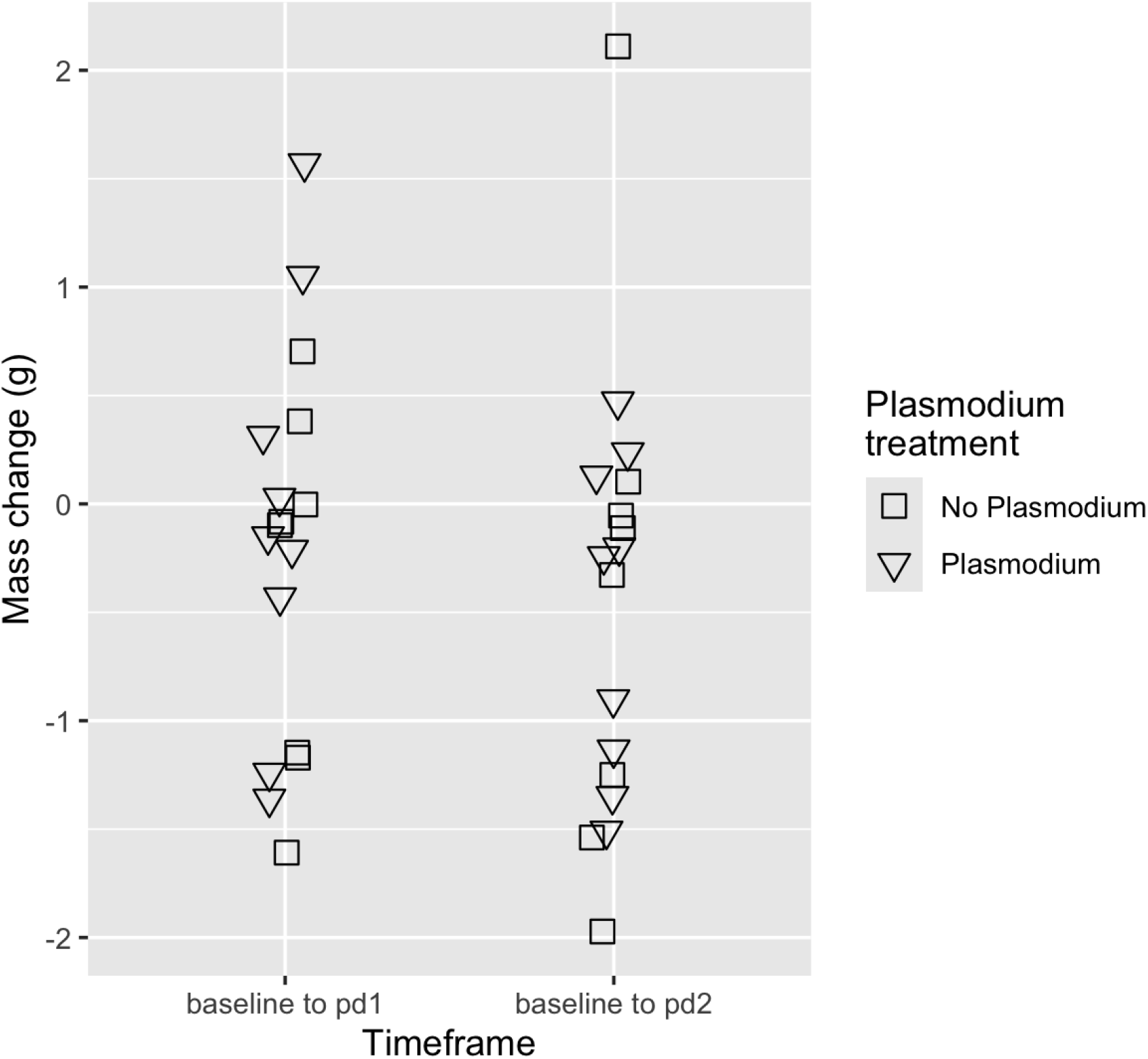
Changes in male dark-eyed juncos (*Junco hyemalis*) mass do not vary by *Plasmodium* inoculation. Data represent changes in mass (g) over two timeframes: from baseline to post-dose one, and baseline to post-dose two. At each dosing point, juncos were given either ivermectin or a carrier control. Points represent data from individual juncos and are jittered for easier viewing.

